# Neural basis of Transcranial Magnetic Stimulation at the single-cell Level

**DOI:** 10.1101/405753

**Authors:** Maria C. Romero, Marco Davare, Marcelo Armendariz, Peter Janssen

**Author notes:** These authors are joint first authors on this work (Maria C. Romero, Marco Davare).

## Abstract

Transcranial Magnetic Stimulation (TMS) can non-invasively modulate neural activity in humans. Despite three decades of research, the spatial extent of the cortical area activated by TMS is still controversial. Moreover, how TMS interacts with task-related activity during motor behavior is unknown. We applied single-pulse TMS over the macaque parietal cortex while recording single-unit activity at various distances from the center of stimulation during grasping. The spatial extent of the TMS-induced activation was remarkably restricted, affecting single neurons in a volume of cortex measuring less than 2 mm. In task-related neurons, TMS evoked a transient excitation followed by reduced activity, which was paralleled by a significantly longer grasping time. Furthermore, TMS-induced activity and task-related activity did not summate in single neurons. These results furnish crucial experimental evidence for the neural basis of the TMS effect at the single-cell level and uncover, the neural underpinnings of behavioral effects of TMS.

## Introduction

Presently, a host of noninvasive brain stimulation (NIBS) techniques are widely used in neuroscience research (Wagner et al., 2007; Bestmann and Feredoes, 2013; Dayan et al., 2013; Miniussi et al., 2013). The most attractive feature of these NIBS techniques is that they allow excitation or inhibition of neural tissue through the skull, so that causal inferences about the role of cortical areas can be made in healthy volunteers. Of all NIBS techniques, Transcranial Magnetic Stimulation (TMS) has been successfully and widely used in volunteers and patients for over thirty years (Hallett, 2000; Pascual Leone et al., 2000; Walsh and Cowey, 2000; Gothe et al., 2002; Rossini and Rossi, 2007; Ziemann, 2011), but a number of fundamental questions remain unresolved. We know very little about the spread of the induced currents in the brain and their excitatory or inhibitory effects on neural activity at the single-cell level, both locally (i.e immediately below the TMS coil) and at a distance (i.e. the response in interconnected brain areas). Inferences about the causal role of a cortical area in behavior critically depend on the ability to manipulate neural activity with high spatial selectivity, since interfering with multiple areas simultaneously could lead to erroneous conclusions. Selective spatial targeting also involves focusing on specific neuronal populations within a cortical area, but again, direct single-cell evidence for the effect of TMS on different neuronal subpopulations is lacking. Furthermore, numerous studies have demonstrated that the application of TMS can cause behavioral effects (Takeuchi et al., 2005; Tunik et al., 2005; Kim et al., 2006; Breitmeyer et al., 2004; Davare et al., 2006; Davare et al., 2010; Reichenbach et al., 2011), yet how TMS-induced activity interacts with ongoing neural activity during behavior is entirely unknown.

Previous studies have employed computational modeling (Opitz et al., 2011; Pashut et al., 2011; De Geeter et al., 2015; Krieg et al., 2015; Thielscher et al., 2015) or indirect measurements of the effect of TMS on neural activity, including behavior (the Motor Evoked Potential or MEP: Muellbacher et al, 2000; Touge et al., 2001; Tsuji and Rothwell, 2002; Fischer and Orth, 2011; Tsang et al, 2014), noninvasive electrophysiology (EEG: Ilmoniemi et al., 1997; Massimini et al., 2005; Thut et al., 2005; Ives et al., 2006) or functional imaging (Bohning et al., 1998; Baudewig et al., 2001; Bestmann et al., 2004; Denslow et al., 2005; Ruff et al., 2006, 2008, Bergmann et al., 2016). Other studies have investigated the effect of repetitive TMS (rTMS) on hemodynamic, local field potential (ECoG) and single-cell responses in anesthetized animals (Allen et al., 2007; de Labra et al., 2007; Pasley et al., 2009; Papazachariadis et al., 2014; Kim et al., 2015). However, none of these approaches provide the spatial and temporal resolution necessary to examine the impact of single-pulse TMS on individual neurons, the spatial extent of TMS effects, nor the important interaction between TMS-induced and task-related activity during behavior. Recently, Mueller et al. (2014) demonstrated the feasibility of measuring single-cell activity during single-pulse TMS, and Ortuño et al. (2014) recorded extracellular activity in remote subcortical brain structures in awake behaving monkeys immediately after repetitive TMS. The monkey model offers several important advantages for TMS studies because of its brain size, the pattern of sulci and gyri (which affects the current spread in the brain, Opitz et al., 2011) comparable to that of the human brain, and the possibility to measure single-cell activity during complex visuomotor tasks (Gu and Corneil, 2014).

## Results

### Modeling of the electric field induced by TMS

To study the TMS influence on neuronal activity, we performed model-based simulations of the induced electrical field and conducted combined single-pulse TMS and extracellular recordings in parietal area PFG during visually-guided grasping (VGG). First, we modelled the spatial spread of the TMS induced current TMS effect over the macaque parietal cortex using existing software (Figure 1B, simNIBS, Opitz et al., 2011), simulating a distance of 15 mm between our TMS coil and the cortical surface (consistent with MR and CT imaging of both animals). According to the simNIBS model, TMS should induce a widespread activation in parietal cortex, extending to frontal and occipital cortex (Figure 1B, left panel). In our single-cell experiments, we applied TMS at 120% of the resting Motor Threshold (rMT), which corresponds to the maximal electric field value Fig 1B (see online methods). To quantify the simulated TMS spread at this intensity (120% rMT), we computed the electric field value associated with the smallest effective TMS intensity capable of recruiting neural tissue (i.e. 100% rMT, corresponding to 83% of the maximal electric field under the coil). Therefore, the simulated spread of a TMS pulse at 120% of the rMT is comprised between 83-100% electric field values (see Thielscher et al., 2015). With this procedure, the width of the spread measured approximately 20 mm both in the anterior-posterior and medio-lateral direction, which is similar to the presumed spread in humans (Anand and Hotson, 2002). On a coronal rendering of the macaque brain (right panel in Figure 1B), the simulation predicts an activation that is mainly limited to the cortex of the parietal convexity, excluding the region buried in the intraparietal sulcus. Next, we measured the spiking activity of neurons located under the center of the coil (7 mm in the antero-posterior axis; 3 mm in the medio-lateral axis), and analyzed the actual spatial spread of TMS applied at different task epochs (light onset -the visual phase of the VGG task- and hand lift - the reaching phase of the VGG task). Also, to verify that the evoked effect was a direct consequence of neuromodulation, we applied TMS at two different intensities: 60% (low intensity TMS) and 120% (high intensity TMS) of the rMT. Using this protocol, we recorded the activity of 538 parietal neurons (476 in the standard experiment and 62 in a control experiment; M1: 172, M2: 304) in two rhesus monkeys. To record as closely as possible to the skull, we precisely positioned the TMS coil as to target the parietal convexity (area PFG, Rozzi et al., 2008) based on anatomical MRI and CT imaging (Figure 1A). Single-pulse TMS induced an artifact in the recordings lasting between 8 and 12 ms (Supplementary Figure 2). Therefore, we excluded the epoch between 0 and 10 ms after TMS onset for all analyses.

**Figure 1.**
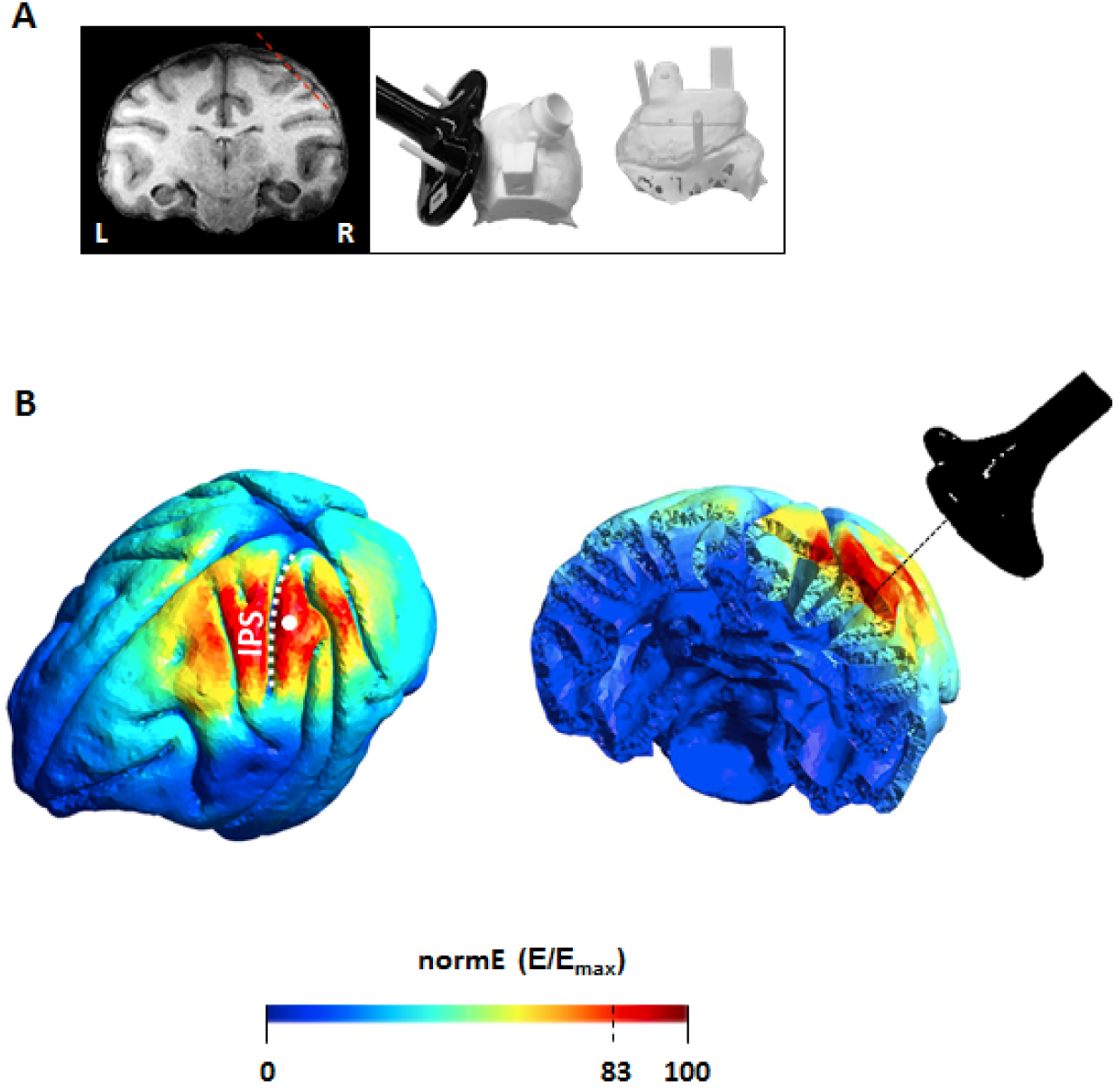
Brain targeting and model of the spatial spread. **A:** Anatomical magnetic resonance image (left) and 3D models of the monkey’s skull (right) indicating respectively, the trajectory of the electrode during recordings (red, dashed line) and the coil positioning method. The right panel shows a top view (left) and a lateral view (right) of a 3D model. During the experiments, a 25 mm figure-of-eight TMS coil (black) was rigidly anchored to the monkey’s skull to allow precise and reproducible coil positioning across recording sessions. Two guiding rods were attached to the monkey’s head implant based on MRI estimations of the cortical target coordinates. B: TMS-induced electric field as modelled with simNIBS. Left: rotated model of the brain indicating the normalized electric field distribution (spatial spread) calculated for the D25 coil. The white dot indicates the center of stimulation (center of the coil). Right: Coronal section of the brain across PFG.

### TMS effect on single neurons

Single-pulse TMS evoked a variety of effects on individual neurons in parietal cortex, as illustrated in Figure 2. By far the most frequently encountered TMS effect was a short-latency burst of action potentials, starting in the first time bin after the TMS artifact (10 ms after the TMS pulse) and lasting less than 50 ms (Figure 2A, left panels). The raster plot of the example neuron in the light-onset condition (top row) in Figure 2A clearly illustrates that in the 8 stimulation trials randomly interleaved with no-stimulation trials, TMS elicited a virtually identical burst of action potentials in the first 30 ms after TMS onset, with the first evoked spikes detected as early as 11 ms after TMS onset. In the first three time bins (10-40 ms after TMS onset), the activity after high-intensity stimulation was significantly higher than in the no-stimulation condition (bin-by-bin analysis, two-sided Wilcoxon ranksum test; p < 0.01), whereas low intensity stimulation did not induce any effect (Figure 2A, right panels). The example neuron in Figure 2A also illustrates that TMS applied in different time epochs of the task frequently induced similar responses, since the TMS-evoked response was virtually identical whether we applied TMS at light onset (top row) or at hand lift (bottom row in Figure 2A, ANOVA, interaction between stimulation-no stimulation and trial epoch, F = 0.04, p = 0.84; df =1). Single-pulse TMS did not exclusively induce excitatory effects in single neurons. The second example neuron (Figure 2B) showed a complex pattern of TMS-induced activity. After an initial short-latency excitation (significantly elevated activity in the epoch 10-40 ms after TMS onset), the neuron became temporarily inhibited for approximately 50 ms, after which a second excitatory phase emerged lasting until 250 ms after TMS onset. This excitation–inhibition–excitation pattern was observed when TMS was applied in either trial epoch (ANOVA, interaction between stimulation – no stimulation and trial epoch, F = 0.27, p = 0.60; df = 1). Overall, the example neuron in Figure 2B clearly demonstrates that the application of TMS in awake behaving animals can induce both excitatory and inhibitory effects, which can reverberate through the cortical network for several hundreds of milliseconds. These two example neurons in Figure 2 did not show task-related activity, but single-pulse TMS induced very similar effects in parietal neurons with task-related activity (see below, Figure 4).

**Figure 2.**
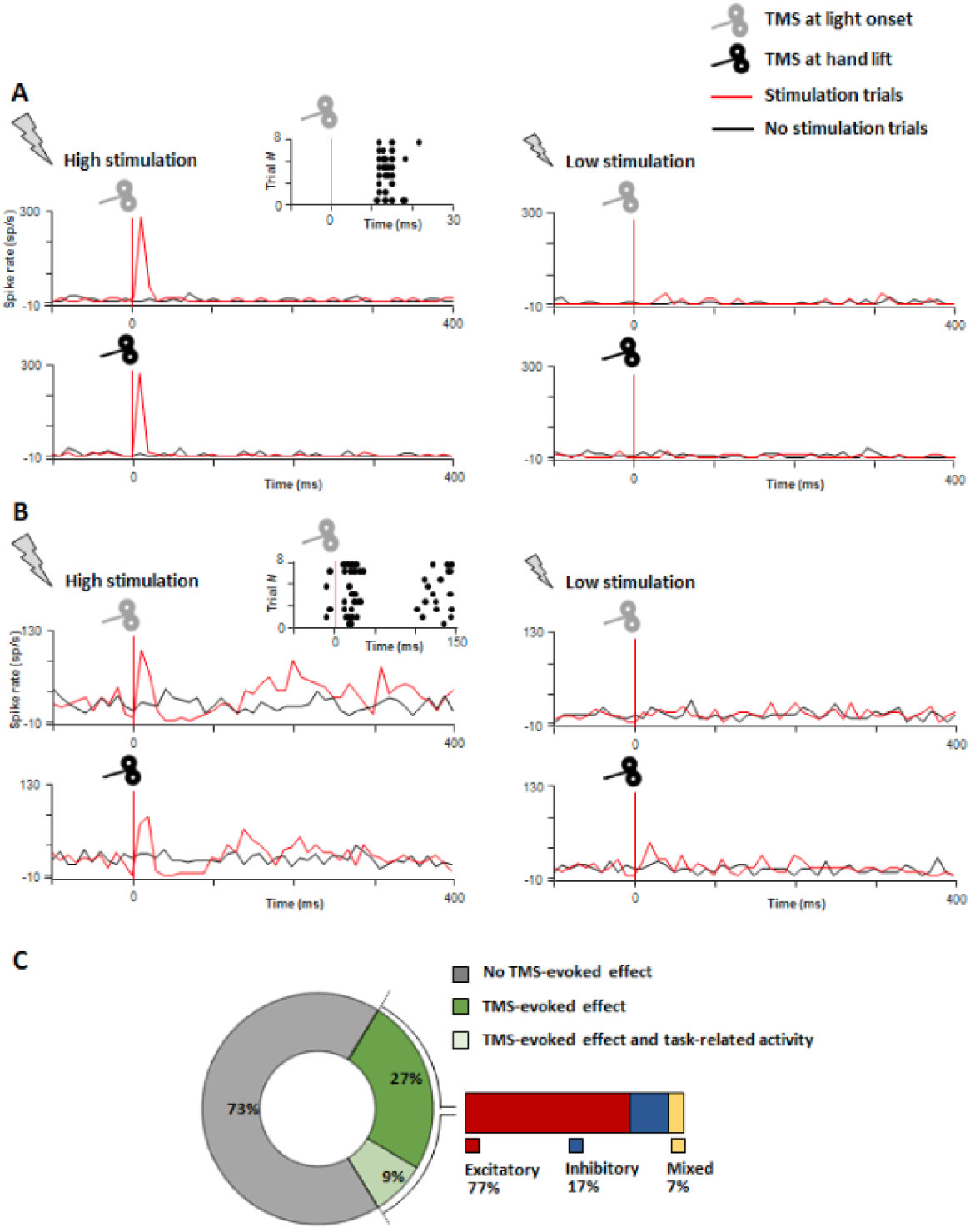
Combined single-pulse TMS and extracellular recordings in parietal area PFG. **A:** Example neuron exhibiting a short-lasting excitatory response to high TMS stimulation (120% rMT; left panel) compared to low stimulation (60% of rMT; right panel). Stimulation (red line plots) and no stimulation trials (black line plots) were randomly interleaved during the task. In a vast majority of PFG neurons, single-pulse TMS evoked a short-lasting burst of activity, which emerged around 20 ms after stimulation, and lasted for approximately 40 ms. The induced excitation did not depend on the time of the stimulation, reaching similar amplitudes when the pulse was applied at ‘light’ onset (grey coil: visual epoch of the VGG task) or at ‘hand lift’ (black coil: reaching epoch of the VGG task). A detailed, trial-by-trial, illustration of the excitatory effect is shown in the raster plot. B: Second example neuron exhibiting a combined excitatory-inhibitory pattern of TMS-evoked activity in response to high (left panel) but not low stimulation (right panel). After an initial excitatory response, we observed a longer-lasting neuromodulation phase combining both inhibitory and excitatory periods for at least 300 ms. Similarly to A, the TMS-induced excitation did not depend on whether TMS was applied during the visual (grey coil) or reaching (black coil) epoch of the task. C: Proportion of PFG recorded neurons with no TMS-evoked effect (grey), TMS-evoked effects (green) and both TMS-evoked and task-related neural responses (light green). The inset on the right shows that amongst cells with TMS-evoked effects, we predominantly found excitatory (red), and few inhibitory (blue) or mixed (yellow) TMS-induced neural responses.

### Spatial extent of the TMS effect

Out of 476 neurons recorded in the standard experiment in two monkeys, 132 (28%) were significantly affected by single-pulse TMS (Figure 2C), i.e. showed at least one time bin with a significant difference between the TMS and the no-TMS conditions at p < 0.01 (two-sided Wilcoxon ranksum test; 35% in monkey Y, 23% in monkey P). Of those 132 neurons, a large majority (77%) showed facilitation (72% in monkey Y, 79% in monkey P), a smaller fraction (17%) showed inhibition (monkey Y: 16%, monkey P: 18%), and an even smaller fraction (7%) showed both facilitation and inhibition (monkey Y: 11%, monkey P: 3%; Figure 2C). In a small fraction of the neurons (11%; monkey Y: 11%, monkey P: 10%), the magnitude of the TMS-induced response depended on the time of stimulation (ANOVA, significant interaction between the factors epoch and *stimulation,* p < 0.05). The example neurons in Figure 2A and B were recorded in the parietal convexity at the center of stimulation, immediately under the TMS coil (approximately 15 mm under the skull), where the TMS-evoked response was maximal. However, to chart the spatial extent of the TMS effect, we also recorded at different distances from the center of stimulation with a spacing of 1 mm, both in the anterior-posterior direction (7 grid positions in monkey Y; 9 in monkey P) and in the medio-lateral direction (3 grid positions in both monkeys) (Figure 3A). The average population responses recorded at the 5 central positions in the grid (Figure 3B and C) reveal the unexpected focality of single-pulse TMS effects. In monkey Y, two recording sites spaced a mere 1 mm apart showed a clear and transient increase in activity (two-sided Wilcoxon ranksum test comparing high stimulation and no stimulation trials; p = 0.019 and 0.014), whereas the surrounding recording sites were barely affected by TMS (Figure 3B). The recording site located 1 mm posterior to the center of stimulation also showed a weaker but significant effect (two-sided Wilcoxon ranksum test; p = 0.016). In the second monkey (Figure 3C), we observed the main TMS-induced response in a single recording site (two-sided Wilcoxon ranksum test; p = 0.006), which was surrounded by recording sites showing no effect of TMS. The interaction between grid position and TMS was significant in a nested ANOVA analysis (with cells as a nested variable) in both animals (p = 0.0002 in monkey Y and p = 0.0001 in monkey P). In addition, we observed a second marginally significant (two-sided Wilcoxon ranksum test; p = 0.038) focus 3 mm away from the center (Supplementary Figures 3 and 4). Figure 3B and C also shows that low intensity single-pulse TMS had very weak effects in general, although we could detect excitatory effects of low-intensity TMS in individual neurons. The average population activity also illustrates that the TMS-evoked response was confined to the first 100 ms after TMS onset, although individual neurons occasionally showed TMS effects lasting up to 200 ms (Supplementary Figures 3, 4 and 6).

**Figure 3.**
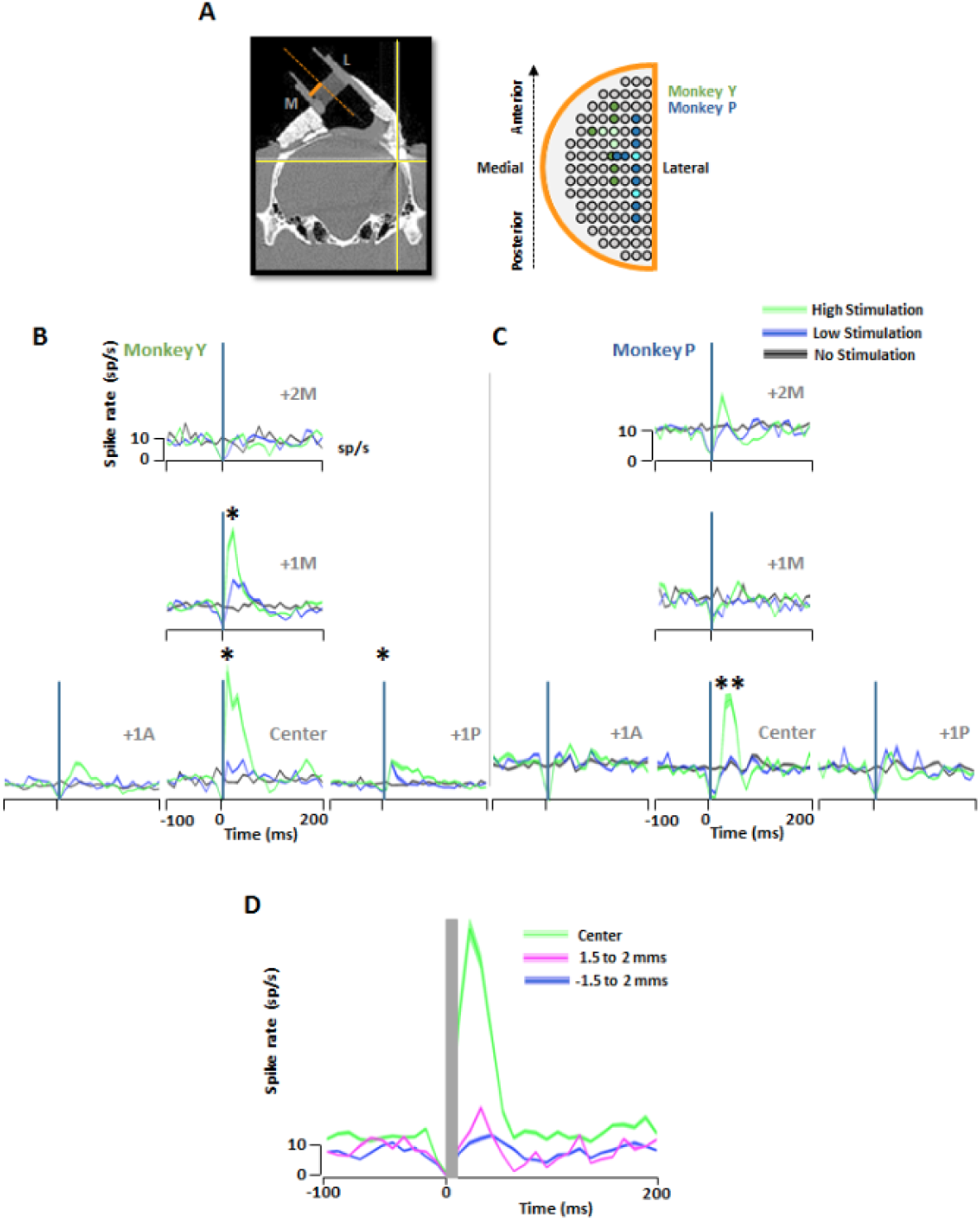
Spatial spread of the TMS-evoked activity. **A:** Diagram of the recording grid (orange) used during the experiments. Colored electrode positions in the grid (green: monkey Y; blue: monkey P) indicate the extent of the area mapped in each animal. Brighter color shadings indicates grid points with significant TMS-evoked effects (i.e. hotspots). **B-C:** Average spike rate obtained for all neurons recorded per grid position in response to high stimulation (green: 120% rMT), low stimulation (blue; 60% rMT) and no stimulation (black), when TMS was applied at ‘light’ onset. Graphs are arranged in a T-shape fashion in order to represent the activity maps spatially for grid positions located immediately under the center of the TMS coil (area mapped: from center to ± 1mm antero-posterior and +2 mm medial). Asterisks indicate statistical strength (*= p ≤ 0.05; **= p ≤ 0.01). B, Monkey Y and C, Monkey P. D: Analysis of the spatial spread along the electrode penetration axis (dorso-ventrally and medio-laterally oriented). The CT scan (A, left panel) illustrates the trajectory of the electrode (orange dotted line). The line plots show the average spike rate observed for all neurons recorded at the same central grid position (aligned to the center of the coil) but at different electrode depths (green: 0 depth or center of stimulation; blue: 1.5 to 2 mms dorsal to the center of stimulation; magenta: 1.5 to 2 mms ventral to the center of stimulation) in response to high TMS stimulation (120% rMT). All averaged data were obtained for the TMS applied at ‘light’ onset.

We based the above analyses and plots on a compilation of all neurons recorded per grid position. However, we also noticed a high degree of spatial selectivity in the TMS-evoked response along the electrode track, i.e. in the dorso-ventral direction, occurring around the center of stimulation. To illustrate this observation, we first identified the depth at which the TMS-evoked response was maximal for every electrode penetration (considered as the center of stimulation; Figure 3B and C). Next, we averaged the activity for all neurons recorded within 500 microns of this position (the center depth) and all neurons recorded 1.5-2 mm dorsal and 1.5-2 mm ventral to this position (Figure 3D). The analysis of the dorso-ventral spread was possible in 7 sessions in monkey Y (35 cells) and 5 sessions in monkey P (36 cells). While the TMS-induced response at the center depth was very robust (N = 25 neurons, 64% showing a significant TMS effect), single-pulse TMS evoked barely any response at the neighboring depths. Our results estimate the spatial selectivity of the TMS effect as a volume of cortex measuring less than 2 mm on a side, at least one order of magnitude smaller than the simulation results (Figure 1B). Figure 3D also illustrates that the magnitude of the TMS-induced response is effectively underestimated in the plots of Figure 3B and C, since the latter contain neurons recorded at different depths. In order to visualize both the average TMS-evoked responses and the variability across neurons, we plotted the net evoked activity of every cell recorded at 9 grid positions (0-3 mm anterior, posterior and medial to the center of stimulation) when we applied TMS at light onset (Supplementary Figure 3). Note that not every neuron in the center was activated by single-pulse TMS. Neurons with a significant TMS effect showed an average transient increase in activity of 27 spikes/sec in the interval from 10 to 80 ms after TMS onset.

Our recordings consisted mainly of very large action potentials that could be recorded for at least one hour and with an average signal-to-noise ratio of 3 or more, as in the example recording in Supplementary Figure 2. However, we also observed significant and virtually identical TMS-induced responses in multi-unit recordings (signal-to-noise ratio of 3 or less; see Supplementary Figure 5A and B). Thus, single-pulse TMS might activate many different neuron types in parietal cortex.

### TMS effect on task-related activity

We also examined how TMS-evoked activity interacted with task-related activity in parietal cortex. In total, 41 neurons (16 in monkey Y and 25 in monkey P) showed significant TMS-induced responses (in the interval between 10 and 200 ms after TMS onset) and significant task-related activity (in the interval between 10 and 400 ms after the hand lift). The example neuron (Figure 4A) and the normalized average activity plot (Figure 4B top panel) illustrate that the TMS-induced response was strong but limited to the first 100 ms after the hand lift, while task-related activity only emerged 200 ms after hand lift. Interestingly, the average normalized activity in TMS trials was significantly lower compared to no-TMS trials in the interval 300-700 ms after hand lift (two-sided Wilcoxon ranksum test; p = 0.047), which was not the case in the low stimulation trials (p = 0.453, Figure 4B lower panel). In total, 11 out of 41 neurons (27%) showed a significant reduction in task-related activity after TMS, which was on average 67% of the task-related activity in the absence of TMS. Thus, single-pulse TMS caused an initial burst of action potentials followed by a prolonged reduction in PFG activity during object grasping. Since task-related activity appeared relatively late in the trial (in the later part of the transport phase of the hand and around the lift of the object) and the TMS-evoked response was short, we could not determine whether task-related activity and TMS-induced activity would linearly summate in PFG neurons. Therefore, we ran a control experiment in which we applied single-pulse TMS 400 ms after the hand lift, at a moment when the average task-related activity was high (N=62). To assess the extent to which TMS-evoked activity and task-related activity summate, we aligned the activity to the onset of the TMS pulse, and compared the average activity in trials when TMS occurred at hand lift (TMS at lift trials) to the activity when TMS occurred 400 ms after hand lift (TMS at lift+400 trials, Figure 4C).

**Figure 4.**
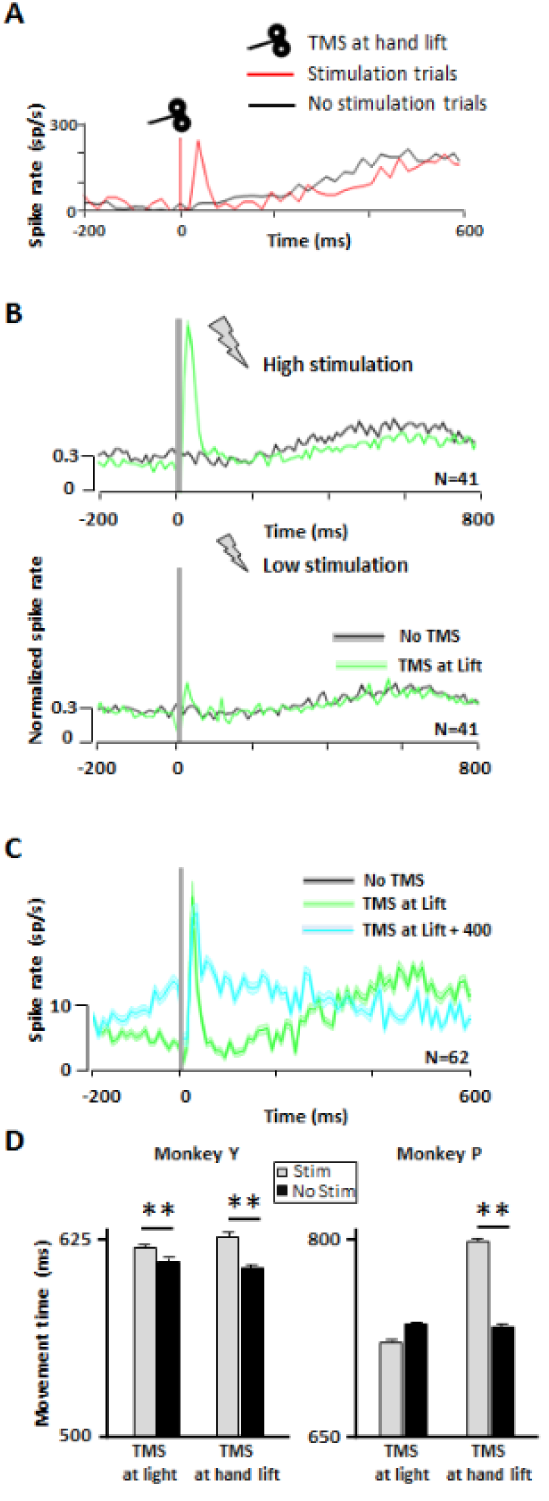
Interaction between TMS effects, task-related activity and behavior. **A:** Example neuron showing combined TMS and task-related activity (same conventions as in Figure 2). The TMS-induced effect was, on average, similar to that described for non-task-related neurons, and did not overlap with the task-related activity. However, the TMS-induced excitation affected the total responsiveness of the neuron, which showed a global decrease in the firing rate during the reaching epoch (250 to 400 ms). B: Normalized spike rate collected during the VGG task comparing the average activity observed at the center of stimulation for all cells showing both significant TMS-evoked effects and task-related activity in the two animals. All neurons were aligned at hand lift and compared across TMS intensities: high (green) versus no stimulation (black; higher panel) and low (green) versus no stimulation (black; lower panel). C: Average spike rate obtained for all task-related neurons recorded in monkey P when TMS was applied at different timings during the reaching epoch (green: TMS at ‘hand lift’; cyan: TMS at ‘hand lift+400ms’). The TMS-induced activity peak was similar across conditions. D: Behavioral effect of TMS. The bar graphs show the total grasping movement times (time elapsed between the hand lift and object pulling) for both monkeys during TMS (grey bars) and no TMS trials (black bars). The effects of 2 different TMS timings are shown: TMS at ‘light’ onset (visual epoch) and TMS at hand lift (reaching epoch). TMS applied during the reaching epoch significantly increased the grasping movement time in both monkeys. Asterisks indicate statistical strength (**= p ≤ 0.01).

Before the TMS pulse, the activity was higher in TMS at hand lift+400 trials than in TMS at hand lift trials because task-related activity was stronger late in the trial. Conspicuously, the TMS-evoked response in TMS at hand lift+400 trials peaked around 40 ms after TMS onset but remained significantly lower compared to that observed at hand lift trials (two-sided Wilcoxon ranksum test; p = 0.022, Wilcoxon ranksum test), indicating that the TMS-evoked response did not summate with task-related activity. Later in the trial (40 – 260 ms after TMS onset), the activity difference reappeared (two-sided Wilcoxon ranksum test; p < 0.001). These observations strongly suggest that TMS-induced activity and task-related activity do not linearly summate in PFG neurons. Importantly, while the above results show that single-pulse TMS interfered with task-related activity in parietal neurons (Figure 4), we also found that single-pulse TMS caused significant behavioral effects in both animals. High-intensity TMS applied immediately after the hand lift significantly increased the time to grasp the object in both monkeys (Figure 4D; a similar lengthening of the total grasp time was present when TMS occurred after light onset, but only in monkey Y). We did not observe any TMS effect on reaction times, and low-intensity TMS had no effect on behavior (Figure 4D). Thus, the prolonged reduction in neural activity in PFG after TMS onset was paralleled by an increase in the total grasp time.

## Discussion

Single-pulse TMS induced a highly localized and short-lived excitation of single neurons in parietal cortex of rhesus monkeys, which interfered with normal task-related activity. These findings have major implications for the interpretation of a very large number of TMS studies in human volunteers and patients.

The spatial selectivity of the TMS effect was unexpected based on previous studies modeling the electric field induced by TMS in humans (e.g. Opitz et al., 2011). We also simulated the electric field induced by TMS in the macaque brain based on state-of-the-art software (simNIBS), and found a predicted spread of at least 10 times broader compared to our single-cell results. However, TMS of primary motor cortex can elicit individual finger movements in humans (Gentner and Classen, 2006) and in monkeys (our own observations). Near-threshold intracortical microstimulation studies in monkeys (Nudo et al., 1996) estimate that the area in primary motor cortex for finger flexion and extension measures approximately 4 mm^2^, which is remarkably consistent with our single-cell results. Thus, the seemingly extraordinary focality of the TMS effect on individual neurons is in fact entirely consistent with previous behavioral results obtained with single-pulse TMS. Our data also confirm the limited penetration of TMS with a figure-of-eight coil. We measured robust TMS-evoked responses in the parietal convexity close to the skull, but very little if any effect further from the TMS coil, in more medial recording positions. The exquisite temporal resolution of single-cell recordings revealed that TMS induces a variety of effects in individual neurons in awake, behaving animals. While most neurons become activated within 40 ms of TMS onset, combinations of excitation and inhibition (the latter most likely caused by the activation of neighboring inhibitory interneurons slightly later in time) were not uncommon in our data set. We even observed secondary activations in recording sites further away from the center of stimulation. All these observations corroborate the idea that single-pulse TMS exerts robust effects on networks of neurons, which cannot be appreciated with population measurements such as fMRI, EEG or behavioral measurements.

We measured the effect of single-pulse TMS on the activity of single neurons generating large action potentials that we could record for an extended period of time (up to an hour) during an active motor task. These spikes most likely originated from large to medium-sized pyramidal neurons typical of area PFG (Rozzi et al., 2008). However, TMS also influenced multi-unit activity characterized by small spikes, and – given the transient inhibition periods we observed in neurons with large spikes – also of small inhibitory interneurons. In contrast, even at the center of stimulation, not every neuron recorded showed an effect of single-pulse TMS. This heterogeneity in the susceptibility of individual neurons to TMS pulses may be related to differences in the local orientation of neurons with respect to the induced electric field (Opitz et al., 2011; Thielscher et al., 2011; Bijsterbosch et al., 2012), which is difficult to estimate with our electrode approach. Thus, TMS influences a wide variety of cell types and evokes a variety of effects at the center of stimulation, the sum of which is a transient excitation lasting less than 100 ms.

The effect of single-pulse TMS on the spiking activity of individual neurons we observed in awake behaving monkeys differed markedly from previous observations in anesthetized animals (Moliadze et al., 2003; Li et al., 2017). Notably, we did not observe long-lasting excitation (up to 300 ms after TMS onset) or suppression (up to two seconds after TMS onset), as reported by Moliadze and colleagues (Moliadze et al., 2003). Undoubtedly, the absence of anesthesia and the active engagement in a motor task in our study could explain these differences. The active engagement in a task is also crucial for the similarity of our study to TMS studies carried out in human volunteers (e.g. Leib et al., 2016; Davare et al., 2012).

A unique advantage of our approach is that we could investigate the relationship between TMS-induced activity, task-related activity and behavior. TMS applied after the onset of the hand movement significantly prolonged the total object grasping time in both monkeys. Previous studies have demonstrated that neurons in area PFG are frequently active during object grasping (Rozzi et al., 2008) and during grasping observation (Fogassi et al., 2005), but no reversible inactivation of PFG during grasping has been published. In our experiments, task-related activity in PFG during the hand movement was suppressed after high-intensity single-pulse TMS. The increase in grasping time we observed may arise from this decreased task-related activity in PFG. Alternatively, the fast burst of action potentials immediately after TMS onset may have interfered with task-related activity in downstream cortical areas involved in grasping such as ventral premotor cortex, with which PFG is connected (Premereur et al., 2015). However, applying TMS at 400 ms after the hand lift (i.e. before the object was lifted) did not affect the grasping time, suggesting that the suppression of task-related activity following TMS may have been the most important factor. Our behavioral data strongly resemble previous findings in humans. TMS applied over the anterior IPS in human volunteers disrupts grasping in the presence (Tunik et al., 2005) or absence (Rice et al., 2006) of a perturbation in object orientation requiring online adjustments. Additionally, TMS applied over the anterior IPS also prolongs reaching times when subjects have to correct for visual perturbations during the reaching movement (Reichenbach et al., 2011). Because the position of the TMS coil was fixed in our experiments, we could not stimulate control sites elsewhere in parietal cortex, but the single-cell data clearly showed that single-pulse TMS activated a patch of cortex in area PFG in the parietal convexity. Thus, our study also represents the first causal evidence for area PFG playing a role in visually-guided grasping. Single-pulse TMS during task-related activity could not excite the neurons above their normal activity levels in the absence of TMS, indicating sublinear summation. This result is puzzling, since the average activity in our population of neurons may not have been maximal. Future studies will have to investigate the relationship between the neuronal ‘state’ and its excitability by TMS.

For several decades, noninvasive brain stimulation techniques have proven to be major assets for both systems neuroscience (Hallett, 2007; Sandrini et al., 2011; Dayan et al., 2013) and the clinical practice (clinical applications in patients; Padberg and George, 2009; Corti et al., 2012; Liew et al., 2014). Invasive animal studies combined with behavioral measurements can uncover new and unprecedented insights into the effects of brain stimulation on individual neurons and on behavior, and will undoubtedly contribute to the refinement and development of novel stimulation protocols (Papazachariadis et al., 2014; Hannah and Rothwell, 2017).

## Methods

### Subjects and surgical procedures

Two male rhesus monkeys (Macaca mulatta; monkey Y, 12 kg; monkey P, 10 kg) were trained to sit in a primate chair. A head post (Crist Instruments) was then implanted on the skull with ceramic screws and dental acrylic. For this and all other surgical protocols, monkeys were kept under propofol anesthesia (10mg/kg/h) and strict aseptic conditions. All experimental procedures were performed in accordance with the National Institutes of Health Guide for the Care and Use of Laboratory Animals and the EU Directive 2010/63/EU, and were approved by the Ethical Committee at KU Leuven. Intensive training in passive fixation and visually-guided grasping began after 6 weeks of recovery. Once the monkeys had achieved an adequate level of performance, a craniotomy was made, guided by anatomical Magnetic Resonance Imaging (MRI), over area PFG of the left hemisphere in monkey Y and the right hemisphere in monkey P. The recording chamber was implanted at an angle of approximately 45 deg with respect to the vertical, allowing oblique penetrations into the parietal convexity (Figure 1A). To confirm the recording positions, glass capillaries were filled with a 2% copper sulfate solution and inserted into a recording grid at five different locations while structural MRI (slice thickness: 0.6 mm) was performed. During the experiments, a precise and highly reproducible positioning of the TMS coil was achieved by means of two guiding rods permanently attached to the skull using dental acrylic, oriented at an angle of approximately 90 deg with respect to the recording chamber. To estimate the position of the coil, we used anatomical MRI and Computed Tomography (CT scan) to build 3D printed models of the skull and implant of each animal (Figure 1A, right panel). Thus, we visualized the relative position of the rods on the head, and calculated the distance between the coil and the brain when the coil was positioned over the rods touching the implant. Based on the CT-MR coregistered images, we calculated that the TMS coil was placed approximately 15 mm from the parietal convexity. The coil induced an posterior-anterior (PA) current over PFG.

### Modeling of the electric field induced by TMS in the macaque brain

A tetrahedral head model (mesh file) was created using simNIBS (simulation of Non-invasive Brain Stimulation Methods; Thielscher et al., 2015) and a standard monkey brain template (National Institute of Mental Health Macaque Template: NMT; Seidlitz et al., 2018). The meshes consisted of five different volumes obtained from MRI images (T1; 0.6 mm isotropic resolution) by the progressive segmentation of five tissue types: white matter (WM), grey matter (GM), cerebro-spinal fluid (CSF), skull, and scalp (Thielscher et al., 2015). The assigned conductivity values were fixed: 0.126 S/m (WM), 0.275 S/m (GM), 1.654 S/m (CSF), 0.01 S/m (skull), 0.465 S/m (scalp). Based on those volumes, simNIBS obtained surface reconstructions of the brain. Tissue segmentation was performed in a semi-automatic way using FSL tools (Functional MRI of the Brain Software Library; Jenkinson et al., 2012) which were manually corrected when required. The electric fields were calculated assuming a quasi-static regime (Thielscher et al., 2011; Opitz et al., 2015), according to the following equation:

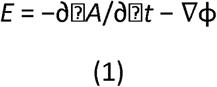

where *E* is the electric field vector and φ denotes the electric potential. The time derivative of the magnetic vector potential *A* of the chosen TMS coil was given as input to the FEM calculations, and the rate of change of the coil current (dI/dt) was set to 1 A/μs. Our model was applied using two different coil files: a Magstim 70 mm figure-of-eight coil (D70) and a Magstim 25 mm figure-of-eight coil (D25; for animal use). The distance between the coil and the cortex was set to 15 mm, as measured in our MRI images, and the coil handle was oriented following the coordinates used in the actual experiment (coil centered on area PFG, projecting along the intraparietal sulcus). The simulations obtained from the two coil types produced similar results, although the total spread was smaller with the smallest coil size (Supplementary Figure 1A). All parameters required for the modelling (change of coil current, distance to the cortex, conductivity values, center of stimulation) were identical across simulations. Using a custom-built measurement coil, we measured the relative output voltage obtained with both the 70 mm and the 25 mm figure-of-eight coils when stimulating at different intensities, taking readings at a 15 mm distance (see Supplementary Figure 1B). Using this procedure, both coils showed similar output intensities (D25 coil 84% of the D70 coil at 70% of the maximum stimulator output intensity). Additionally, we measured the rMT with the two coils (D70 and D25) in two monkeys. The minimum intensity required to evoke finger twitches by applying single-pulse TMS on the contralateral primary motor cortex was very similar using the two coils (D70: ± 40% and D25: ± 35% of the maximum intensity for monkey 1; D70: ± 46 and D25: ± 46% of the maximum intensity for monkey 2).

### TMS stimulation

TMS was applied at 60% and 120% of the rMT (see below for details) by means of a 25 mm figure-of-eight coil. We determined the rMT for each animal as the lowest stimulus intensity at which TMS of the primary motor cortex produced a contralateral finger or wrist movement, (right hemisphere on monkey Y and left hemisphere on monkey P) while the monkey held its hand in the resting position. In monkey Y, we used a Magstim 200 Stimulator (Magstim, UK), which delivered monophasic pulses (100 μs rise time, 1ms duration, 120% rMT = 60% of max stimulator output), whereas in monkey P, we used a Magstim Rapid Stimulator (Magstim, UK), which delivered biphasic pulses (100 μs rise time, 250 μs duration; 120% rMT = 70% of max stimulator output). In a series of control experiments, we also used the Magstim Rapid Stimulator in monkey Y, and could reproduce the basic findings obtained with the Magstim 200. For the duration of the experiment, the coil was placed over the guiding rods tangentially to the skull. To provide additional stability, the TMS coil was held in place by means of an adjustable metal arm. Since our recording sessions averaged 4 hours in duration, gel-filled cool packs were placed around the coil to prevent overheating (Nexcare; 3M Company, Minnesota, USA).

### Single-cell recordings during single-pulse TMS

During the application of single-pulse TMS, we recorded single-unit activity with tungsten microelectrodes (impedance: 1 MΩ at 1 kHz; FHC) inserted through the dura by means of a 23-gauge stainless steel guide tube and a hydraulic microdrive (FHC). Mimicking the artifact reduction strategy proposed by Mueller and colleagues (Mueller et al., 2014), we used diodes and serial low-gain amplification to clip the artifact generated by the magnetic pulses and to prevent amplifier saturation. For this, we modified a regular BAK Electronics preamplifier (Model A-1) by connecting two leakage diodes (BAS45A) anti-parallel between the signal lines and ground before each stage of amplification. The initial front-end of the headstage remained unmodified to maintain the high-input impedance. Also, we covered the tungsten microelectrodes with polyamide tubing (TWPT-0079-30-50; Small Parts) to minimize vibration inside the guide tube. With these settings, the evoked TMS artifact in our signal ranged from 8 to 12 ms (Supplementary Figure 2). Neural activity was amplified and filtered (300-5,000 Hz) following a standard recording protocol for spike detection. Using a dual time-window discriminator (LabVIEW and custom-built software), we isolated individual neurons and the TMS artifact, which was detected online and subtracted from the neural data. In addition, we recorded the entire raw signal (after filtering) for the analysis of multi-unit activity. We monitored the right eye position using an infrared-based camera system (Eye Link II, SR Research, Ontario, Canada) sampling the pupil position at 500 Hz.

### Grasping task

During the experiments, the monkeys were sitting upright with the head fixed. For each recording session, the hand ipsilateral to the recording chamber remained restrained within the chair, while the contralateral hand was placed on a resting device. A single grasping object (large sphere; diameter: 35 mm) was located in front of the monkey at a distance of 30 cm from the resting position. The resting position of the hand, the start of the reach to grasp movement and the pull of the object were detected by fiber-optic cables. Our motor task consisted of a visually guided grasping task (VGG), in which the monkey had to place the hand contralateral to the recorded hemisphere in the resting position in complete darkness to initiate the sequence. After a variable time (intertrial interval: 2000-3000 ms), a red laser projected a fixation spot under the object. If the animal maintained its gaze inside the electronically-defined fixation window (+/- 2.5 deg) for 500 ms, the object was illuminated. Following a variable delay (900-1100 ms), a visual GO cue (dimming of the laser) instructed the monkey to lift the hand from the resting position, and reach, grasp, lift and hold the sphere for a variable interval (holding time, 500-900 ms). Whenever the monkey performed the whole sequence correctly, it received a drop of juice as reward. During the task, we measured both the time between the go-signal and the onset of the hand movement (reaction time), and the time between the start of the movement and the lift of the object (grasping time).

### Experimental protocol

To address the influence of single-pulse TMS on neuronal activity, we interleaved trials with (50% of the trials) and without (50% of the trials) stimulation, and applied single magnetic pulses in different task epochs. These TMS pulses were applied in blocks of either low (60% rMT) or high (120% rMT) intensity TMS, randomly interleaved with no-stimulation trials. In both monkeys, we applied single-pulse TMS at two different task epochs: at light onset above the object, and immediately (within 2 ms) after the onset of the hand movement. In monkey P, we also run an experimental control to investigate the effect of TMS on task-related activity. For this, we applied high and low intensity TMS at both hand lift and at hand lift+400 ms. Across multiple experimental sessions, we measured the TMS-induced changes in the spiking activity of individual neurons located in a region under the center of the coil extending 7 mm in the antero-posterior direction and 3 mm in the medio-lateral direction (Supplementary Figures 3 and 4).

### Data analysis

All data analyses were performed in MATLAB (MathWorks, Massachusetts, USA). For the high- and low stimulation trials, the neural activity was aligned on the TMS pulse. Also, for comparison, the no-stimulation trials were aligned on the same time bin (either light onset or hand lift). Net neural responses were then calculated as the average firing rate recorded after TMS minus the baseline (spike rate calculated from the mean activity of the cell in the 800 ms interval preceding TMS).

We created line and raster plots comparing the average response (spikes/sec) of every cell during no stimulation, low stimulation and high stimulation trials at different task epochs. A bin-by-bin analysis (two-sided Wilxocoxon ranksum test; bin size: 20 ms) comparing the spike rate in high stimulation bins with that in the corresponding no-stimulation bins was performed to classify the neurons according to their functional properties (excitatory TMS effect, inhibitory TMS effect, or combination of excitatory and inhibitory TMS effects). Similarly, we also ran a two-sided Wilxocoxon ranksum test to identify those neurons with significant task-related activity. For this, we first aligned the neural activity on the lift of the hand and then compared the cell’s responses obtained during the delay (−200 to 0 ms before the lifting of the hand) and the reaching-to-grasp phases of the task (10 to 410 ms after lifting the hand/TMS event).

To determine the significance of the TMS-evoked effect in individual cells, we compared the cell responses observed between 10 and 50 ms post-TMS in the high stimulation condition to those in the no-stimulation condition (two-sided Wilxocoxon ranksum test). A two-way ANOVA was performed to quantify the interaction between the factors *stimulation and task epoch*. To estimate the spatial spread of the observed TMS effect, we calculated the average response of all neurons recorded at each recording position during low-, high- and no-stimulation trials. For each recording grid position, we determined the significance of the TMS-evoked effect by performing a Wilcoxon ranksum test comparing the average spike rate across all neurons recorded at that position. In this analysis, we compared the bin showing the maximum response to stimulation (high stimulation trials) to the corresponding bin in the no stimulation trials (bin size: 20 ms). Next, we tested whether the TMS-evoked effect differed significantly between the center of stimulation (i.e., the recording position with the highest TMS effect) and four neighboring positions 1 mm away (+1 mm anterior, +1 mm posterior, +1 mm medial and +2 mm medial to the center) using a three-way nested ANOVA with factors position and TMS condition (high versus no stimulation), and with cells as nested factors. To normalize the activity across all neurons showing task-related activity, we used the normr function in Matlab.

Finally, we measured behavioral effects of TMS by directly comparing the reaction times and grasping times obtained for the high stimulation and no stimulation conditions with single-pulse TMS applied at light onset and at the onset of the hand movement. To adjust for multiple testing, we applied the Bonferroni correction.

---

**Supplementary Figure 1. simNIBS model and coil measurements. A:** TMS-induced electric field as modelled with simNIBS. Rotated models of the brain showing the normalized electric field distribution (spatial spread) calculated for two different coils: D70 (left) and D25 (right). The white dot indicates the center of stimulation (center of the coil). **B**: Output voltage measurements taken with the D70 and the D25 coils at a distance of 15 mm. Using a home-made mini-coil, we took readings of the evoked voltage by applying single-pulses at different intensities (20-80% of the Magstim Rapid).

**Supplementary Figure 2. TMS artifact**. Voltage graph showing the TMS-evoked artifact on a single stimulation trial. By detecting the peaks we were able to isolate individual spikes (indicated in the graph with an orange asterisk) recorded before, during and after TMS. On an average session, the artifact duration lasted about 10 ms. However, the frequency and intensity of the evoked artifact evolved during this period, decaying towards the end, so that in the last phase it was possible to record cellular activity, with spikes overriding the mechanical noise.

**Supplementary Figure 3. Neuronal variability across PFG. A:** Spatial map detailing the TMS-induced activity across grid positions (from 2 mm anterior to 4 mm posterior to the center of stimulation; and 0 to 2 mm medial to the center of stimulation) in monkey Y. For each individual graph, colored lines indicate the net response of a single neuron recorded at a particular position. TMS delivery is indicated by a dashed red line, occurring in this case at ‘light’ onset. B: Spatial map in monkey P. Same conventions as in A. For both animals, some neurons outside the center of stimulation were also responsive to TMS. However, on average, the number of affected neurons decreased considerably for positions located only 1 mm away from the center of the coil.

**Supplementary Figure 4. Complete map of the spatial spread of the TMS effect, average responses.** A: Spatial spread in monkey Y. Average spike rate of all neurons collected at the center of stimulation and neighboring mapped grid positions, extending over 7 mms in the anterior-posterior axis and 3 mms in the mediolateral axis. The figure is arranged in a T-shape to summarize the activity maps obtained for high stimulation (green: 120% rMT), low stimulation (blue; 60% rMT) and no stimulation (black) when TMS was applied at ‘light’ onset. Asterisks indicate statistical strength (*= p ≤ 0.05; **= p ≤ 0.01). B: Spatial spread in monkey P. Same conventions as in A.

**Supplementary Figure 5. Offline spike sorting. A:** Voltage diagram showing the analyses performed on single spikes to differentiate neuronal size (single- and multi-unit activity). For each individual isolated spike (color lines), a baseline (noise) threshold and a peak threshold (signal= 3 × baseline) were assigned to calculate the signal-to-noise ratio (SNR). All neurons showing an SNR equal or higher than 3 were classified as single-units or megaspikes. In contrast, neurons with an SNR lower than 3 were considered small units, which were often associated to multi-unit recordings. B: Average spike rate obtained for all neurons classified as megaspikes (left) and small-units (right), recorded in both animals at the center of stimulation. A direct comparison of the high stimulation (green: 120% of the rMT) and no stimulation (black) conditions in both populations evidenced that the evoked TMS effect was similar across neuronal types (size).

**Supplementary Figure 6. Latency and duration of the TMS effects on neuronal activity. A**: Bin-by-bin analyses of the TMS effect: latency. Distribution of the first bin (bin size=20 ms) showing a significant TMS effect across all positions tested in the two animals (green: monkey Y; blue: monkey P). The first bin analyzed extended from 10 to 30 ms after TMS onset. B: Bin-by-bin analyses of the TMS effect: duration. For the same analyses, distribution of the last significant bin (bin size=20 ms). Same conventions as in A.

## Acknowledgements

This work was supported by Fonds voor Wetenschappelijk Onderzoek Vlaanderen, Odysseus (G.0007.12), and Program Financing (PFV10/008). We would like to thank Axel Thielscher for providing the coil files and Wouter Depuydt for his assistance with the use of simNIBS. We also thank Stijn Verstraeten, Christophe Ulens, Piet Kayenbergh, Gerrit Meulemans, Marc De Paep, Astrid Hermans and Inez Puttemans for their technical contributions, and Steve Raiguel for comments on a previous version of this manuscript.

## References

Allen E.A., Pasley B.N., Duong T., & Freeman R.D. Transcranial magnetic stimulation elicits coupled neural and hemodynamic consequences. Science, 317, 1918–1921 (2007).

Anand S., Hotson J. Transcranial magnetic stimulation: neurophysiological applications and safety. Brain Cogn, 50, 366–386 (2002).

Baudewig J., Siebner H.R., Bestmann S., Tergau F., Tings T., Paulus W. & Frahm J. Functional MRI of cortical activations induced by transcranial magnetic stimulation (TMS). Neuroreport, 12, 3543–3548 (2001).

Bergmann T.O., Karabanov A., Hartwigsen G., Thielscher A. & Siebner H.R. Combining non-invasive transcranial brain stimulation with neuroimaging and electrophysiology: Current approaches and future perspectives. NeuroImage, 140, 4–19 (2016).

Bestmann S., Baudewig J., Siebner H.R., Rothwell J.C. & Frahm J. Functional MRI of the immediate impact of transcranial magnetic stimulation on cortical and subcortical motor circuits. Eur. J. Neurosci., 19, 1950–1962 (2004).

Bestmann S. & Feredoes E. Combined neurostimulation and neuroimaging in cognitive neuroscience: past, present and future. Ann. N. Y. Acad. Sci., 1296, 11–30 (2013).

Bijsterbosch J.D., Barker A.T., Lee K-H. & Woodruff P.W.R. Where does transcranial magnetic stimulation (TMS) stimulate? Modelling of induced field maps for some common cortical and cerebellar targets. Med. Biol. Eng. Comput., 50, 671–81 (2012).

Bohning D.E., Shastri A., Nahas Z., Lorberbaum J.P., Andersen S.W., Dannels W.R., Haxthausen E.U., Vincent D.J. & George M.S. Echoplanar BOLD fMRI of brain activation induced by concurrent transcranial magnetic stimulation. jInvest. Radiol., 33, 336–340 (1998).

Breitmeyer B.G., Ro T. & Ogmen H. A comparison of masking by visual and transcranial magnetic stimulation: implications for the study of conscious and unconscious visual processing. Conscious Cogn., 13, 829–843 (2004).

Corti M., Patten C., & Triggs W. Repetitive transcranial magnetic stimulation of motor cortex after stroke: a focused review. Am. J. Phys. Med. Rehabil., 91, 254, 270 (2012).

Davare M., Andres M., Cosnard G., Thonnard J.L. & Olivier E. Dissociating the role of ventral and dorsal premotor cortex in precision grasping. J. Neurosci., 26, 2260–2268 (2006).

Davare M., Rothwell J.C. & Lemon R.N. Causal connectivity between the human anterior intraparietal area and premotor cortex during grasp. Curr. Biol., 20, 176–181 (2010).

Davare M., Zénon A., Pourtois G., Desmurget M. & Olivier E. Role of the medial part of intraparietal sulcus in implementing movement direction. Cereb. Cortex, 22, 1382–1394 (2012).

Dayan E., Censor N., Buch E.R., Sandrini M. & Cohen L.G. Noninvasive brain stimulation: from physiology to network dynamics and back. Nat. Neurosci., 16, 838–844 (2013).

De Geeter N., Crevecoeur G., Leemans A. & Dupré L. Effective electric fields along realistic DTI-based neural trajectories for modelling the stimulation mechanisms of TMS. Phys. Med. Biol., 60, 453–471 (2015).

de Labra C., Rivadulla C., Grieve K., Marino J., Espinosa N. & Cudeiro J. Changes in visual responses in the feline dLGN: selective thalamic suppression induced by transcranial magnetic stimulation of VCereb. Cortex, 17, 1376–1385 (2007).

Denslow S., Lomarev M., George M.S. & Bohning D.E. Cortical and subcortical brain effects of transcranial magnetic stimulation (TMS)-induced movement: an interleaved TMS/functional magnetic resonance imaging study. Biol. Psychiatry, 57, 752–760 (2005).

Fischer M. & Orth M. Short-latency sensory afferent inhibition: conditioning stimulus intensity, recording site, and effects of 1 Hz repetitive TMS. Brain Stimul., 4, 202–209 (2011).

Fogassi L., Ferrari P.F., Gesierich B., Rozzi S., Chersi F. & Rizzolatti G. Parietal lobe: from action organization to intention understanding. Science, 308, 662–667 (2005).

Gentner R. & Classen J. Modular organization of finger movements by the human central nervous system. Neuron, 52, 731–742 (2006).

Gothe J., Brandt S.A., Irlbacher K., Röricht S., Sabel B.A. & Meyer B.U. Changes in visual cortex excitability in blind subjects as demonstrated by transcranial magnetic stimulation. Brain, 125, 479–490 (2002).

Gu C. & Corneil B.D. Transcranial magnetic stimulation of the prefrontal cortex in awake nonhuman primates evokes a polysynaptic neck muscle response that reflects oculomotor activity at the time of stimulation. J. Neurosci., 34, 14803–14815 (2014).

Hallett M. Transcranial magnetic stimulation and the brain. Nature, 406, 147–150 (2000).

Hallett M. Transcranial magnetic stimulation: a primer. Neuron, 55, 187–199 (2007).

Hannah R., Rothwell J.C. Pulse duration as well as current direction determines the specificity of Transcranial Magnetic Stimulation of motor cortex during contraction. Brain Stimul., 10, 106–115 (2017).

Ilmoniemi R.J., Virtanen J., Ruohonen J., Karhu J., Aronen H.J., Näätänen R. & Katila T. Neuronal responses to magnetic stimulation reveal cortical reactivity and connectivity. Neuroreport, 8, 3537–3540 (1997).

Ives J.R., Rotenberg A., Poma R., Thut G. & Pascual-Leone A. Electroencephalographic recording during transcranial magnetic stimulation in humans and animals. Clin. Neurophysiol., 117, 1870–1875 (2006).

Kim D.H., Georghiou G.E. & Won C. Improved field localization in transcranial magnetic stimulation of the brain with the utilization of a conductive shield plate in the stimulator IEEE. Trans. Biomed. Eng., 53, 720–725 (2006).

Kim T., Allen E.A., Pasley B.N. & Freeman R.D. Transcranial magnetic stimulation changes response selectivity of neurons in the visual cortex. Brain Stimul., 8, 613–623 (2015).

Krieg T.D., Salinas F.S., Narayana S., Fox P.T. & Mogul D.J. Computational and experimental analysis of tms-induced electric field vectors critical to neuronal activation. J. Neural. Eng., 12, 046014 (2015).

Leib R, Mawase F, Karniel A, Donchin O, Rothwell J, Nisky I, Davare M. Stimulation of PPC affects the mapping between motion and force signals for stiffness perception but not motion control. J. Neurosci., 36, 10545–10559 (2016).

Li B., Virtanen J.P., Oeltermann A., Schwarz C., Giese M.A., Ziemann U. & Benali A. Lifting the veil on the dynamics of neuronal activities evoked by transcranial magnetic stimulation. Elife, e30552 (2017).

Liew S-L., Santarnecchi E., Buch E.R. & Cohen L.G. Non-invasive brain stimulation in neurorehabilitation: local and distant effects for motor recovery. Front. Hum. Neurosci., 8, 378 (2014).

Massimini M., Ferrarelli F., Huber R., Esser S.K., Singh H. & Tononi G. Breakdown of cortical effective connectivity during sleep. Science, 309, 2228–2232 (2005).

Miniussi C., Harris J.A. & Ruzzoli M. Modelling non-invasive brain stimulation in cognitive neuroscience. Neurosci. Biobehav. Rev., 37, 1702–1712 (2013).

Moliadze V., Zhao Y., Eysel U. & Funke K. Effect of transcranial magnetic stimulation on single-unit activity in the cat primary visual cortex. J. Physiol., 553, 665–679 (2003).

Muellbacher W., Ziemann U., Boroojerdi B. & Hallett M. Effects of low-frequency transcranial magnetic stimulation on motor excitability and basic motor behavior. Clin. Neurophysiol., 111, 1002– 1007 (2000).

Mueller J.K., Grigsby E.M., Prevosto V., Petraglia F.W. 3rd, Rao H., Deng Z.D., Peterchev A.V., Sommer M.A., Egner T., Platt M.L. & Grill W.M. Simultaneous transcranial magnetic stimulation and single-neuron recording in alert non-human primates. Nat. Neurosci. 17, 1130–1136 (2014).

Nudo R.J., Milliken G.W., Jenkins W.M. & Merzenich M.M. Use-dependent alterations of movement representations in primary motor cortex of adult squirrel monkeys. J. Neurosci., 16, 785–807 (1996).

Opitz A., Paulus W., Will S., Antunes A., Thielscher A. Determinants of the electric field during transcranial direct current stimulation. NeuroImage, 109, 140–150 (2015).

Opitz A., Windhoff M., Heidemann R., Turner R. & Thielscher A. How the brain tissue shapes the electric field induced by transcranial magnetic stimulation. NeuroImage, 58, 849–859 (2011).

Ortuño T., Grieve K.L., Cao R., Cudeiro J. & Rivadulla C. Bursting thalamic responses in awake monkey contribute to visual detection and are modulated by corticofugal feedback. Front. Behav. Neurosci., 30, 8–108 (2014).

Padberg F. & George M.S. Repetitive transcranial magnetic stimulation of the prefrontal cortex in depression. Exp. Neurol., 219, 2–13 (2009).

Papazachariadis O., Dante V., Verschure P.F. Del Giudice P. & Ferraina S. iTBS-induced LTP-like plasticity parallels oscillatory activity changes in the primary sensory and motor areas of macaque monkeys. PLoS One, 10, e112504 (2014).

Pascual-Leone A., Walsh V. & Rothwell J. Transcranial magnetic stimulation in cognitive neuroscience—virtual lesion, chronomotery, and functional connectivity. Curr. Opin. Neurobiol., 10, 232–237 (2000).

Pashut T., Wolfus S., Friedman A., Lavidor M., Bar-Gad I., Yeshurun Y. & Korngreen A. Mechanisms of magnetic stimulation of central nervous system neurons. PLoS Comput Biol, 7, e1002022 (2011).

Pasley B.N., Allen E.A. & Freeman R.D. State-dependent variability of neural responses to transcranial magnetic stimulation if the visual cortex. Neuron, 62, 291–303 (2009).

Premereur E, Van Dromme I.C., Romero M.C., Vanduffel W., & Janssen P. Effective connectivity of depth-structure-selective patches in the lateral bank of the macaque intraparietal sulcus. PLoS Biol., 13, e1002072 (2015)

Reichenbach A., Bresciani J.P., Peer A., Bülthoff H.H. & Thielscher A. Contributions of the PPC to online control of visually guided reaching movements assessed with fMRI-guided TMS. Cereb. Cortex, 21, 1602–1612 (2011).

Rice N.J., Tunik E. & Grafton S.T. The anterior intraparietal sulcus mediates grasp execution, independent of requirement to update: new insights from transcranial magnetic stimulation. J. Neurosci., 26, 8176–8182 (2006).

Rossini P.M. & Rossi S. Transcranial magnetic stimulation: diagnostic, therapeutic, and research potential. Neurology, 68, 484–488 (2007).

Rozzi S., Ferrari P.F., Bonini L., Rizzolatti G., Fogassi L. Functional organization of inferior parietal lobule convexity in the macaque monkey: electrophysiological characterization of motor, sensory and mirror responses and their correlation with cytoarchitectonic areas. Eur. J. Neurosci. 28, 1569–1588 (2008).

Ruff C.C., Blankenburg F., Bjoertomt O., Bestmann S., Freeman E., Haynes J.D., Rees G., Josephs O., Deichmann R. & Driver J. Concurrent TMS-fMRI and psychophysics reveal frontal influences on human retinotopic visual cortex. Curr. Biol., 16, 1479–1488 (2006).

Ruff C.C., Bestmann S., Blankenburg F., Bjoertomt O., Josephs O., Weiskopf N., Deichmann R. & Driver J. Distinct causal influences of parietal versus frontal areas on human visual cortex: evidence from concurrent TMS-fMRI. Cereb. Cortex, 18, 817–827 (2008).

Sandrini M., Umiltà C. & Rusconi E. The use of transcranial magnetic stimulation in cognitive neuroscience: a new synthesis of methodological issues. Neurosci. Biobehav. Rev., 35, 516–536 (2011).

Seidlitz J., Sponheim C., Glen D., Ye F.Q., Saleem K.S., Leopold D.A., Ungerleider L., Messinger A. A population MRI brain template and analysis tools for the macaque. NeuroImage, 170, 121–131 (2018).

Takeuchi N., Chuma T., Matsuo Y., Watanabe I. & Ikoma K. Repetitive transcranial magnetic stimulation of contralesional primary motor cortex improves hand function after stroke. Stroke, 36: 2681–2686 (2005).

Thielscher A., Antunes A. & Saturnino G.B. Field modelling for transcranial magnetic stimulation: a useful tool to understand the physiological effects of TMS? Conf. Proc. IEEE Eng. Med. Biol. Soc., 2015, 222–225 (2015).

Thielscher A., Opitz A. & Windhoff M. Impact of the gyral geometry on the electric field induced by transcranial magnetic stimulation. NeuroImage, 54, 234–243 (2011).

Thut G., Ives J.R., Kampmann F., Pastor M.A. & Pascual-Leone A. A new device and protocol for combining TMS and online recordings of EEG and evoked potentials. J. Neurosci. Methods, 141, 207– 217 (2005).

Touge T., Gerschlager W., Brown P. & Rothwell J.C. Are the after-effects of low-frequency rTMS on motor cortex excitability due to changes in the efficacy of cortical synapses? Clin. Neurophysiol., 112, 2138–2145 (2001).

Tsang P., Jacobs M.F., Lee K.G., Asmussen M.J., Zapallow C.M. & Nelson A.J. Continuous theta-burst stimulation over primary somatosensory cortex modulates short-latency afferent inhibition. Clin. Neurophysiol., 125, 2253–2259 (2014).

Tsuji T. & Rothwell J.C. Long lasting effects of rTMS and associated peripheral sensory input on MEPs, SEPs and transcortical reflex excitability in humans. J. Physiol., 540, 367–376 (2002).

Tunik E., Frey S.H. & Grafton S.T. Virtual lesions of the anterior intraparietal area disrupt goal-dependent on-line adjustments of grasp. Nat. Neurosci., 8, 505–511 (2005).

Wagner T., Valero-Cabre A. & Pascual-Leone A. Noninvasive human brain stimulation. Annu. Rev. Biomed. Eng., 9, 527–565 (2007).

Walsh V. & Cowey A. Transcranial magnetic stimulation and cognitive neuroscience. Nat Rev Neurosci. 1, 73–79 (2000).

Ziemann U. Transcranial magnetic stimulation at the interface with other techniques: a powerful tool for studying the human cortex. Neuroscientist, 17, 368–381 (2011).

